# Comparative Metagenomics of Soda Lakes and the Rumen Reveals Novel Microbial Lineages and Carbohydrate-active Enzyme Diversity

**DOI:** 10.1101/2025.06.14.659726

**Authors:** Oliyad Jeilu, Addis Simachew, Erica M. Hartmann, Erik Alexandersson, Eva Johansson

## Abstract

Microbial communities have evolved diverse strategies to metabolize similar substrates, shaped by environmental pressures and interspecies interactions. Comparative analyses across extreme and host-associated ecosystems are essential for uncovering novel microbial diversity, functional adaptations, and enzymes with industrial relevance. Here, we present a comparative metagenomic analysis of two ecologically distinct environments, highly alkaline-saline soda lakes, and the host-associated rumen to investigate microbial diversity, metabolic potential, and carbohydrate-active enzyme (CAZyme) repertoires. Taxonomic profiling revealed distinct microbial signatures: soda lake microbiomes were dominated by Proteobacteria, particularly *Halomonas*, and *Nitrincola*, while the rumen was enriched in Bacteroidota, Fibrobacteres, and Firmicutes, with *Prevotella* and *Fibrobacter* as key taxa. The soda lake MAGs exhibit greater evolutionary divergence, suggesting the presence of novel microbial lineages. Codon usage profiling revealed distinct patterns of GC3 bias, suggesting divergent translational optimization strategies shaped by environmental constraints. CAZyme profiling uncovered ecosystem-specific glycoside hydrolase repertoires, with the rumen enriched in plant fiber-degrading enzymes and soda lakes exhibiting broader GH diversity adapted to high salinity and alkalinity. These findings highlight distinct microbial and enzymatic strategies for carbon cycling and position soda lakes as promising reservoirs of extremophile-adapted enzymes for biotechnological applications.

## Background

Microbial life thrives in diverse environments, from chemically harsh ecosystems to highly specialized biological niches^1,2^. Soda lakes are unique poly-extreme environments characterized by high alkalinity (typically pH 9–11) and elevated salinity, offering a distinctive ecological niche for diverse microbial communities^3^. These environments, found globally but especially abundant along the East African Rift Valley, harbor microorganisms capable of surviving and thriving under extreme conditions^4,5^. Ethiopian soda lakes, including Lakes Abijata, Chitu, and Shala, are renowned for their high primary productivity and diverse microbial communities, encompassing bacteria, archaea, and eukaryotes^4,6^.

Parallel to these extreme but productive soda lake ecosystems, another biologically dense and metabolically dynamic microbial habitat exists: the rumen of ruminants. The rumen is a specialized digestive organ in ruminants like cattle, goats, and sheep, where microbial fermentation breaks down plant biomass into usable nutrients for the host animal^7^. The rumen is a warm (∼39–40°C), slightly acidic (pH ∼5.5–6.8), and anaerobic environment. The rumen microbiome is one of the most complex among the wide array of microbial ecosystems present on Earth^8,9^.

Despite their environmental differences, soda lakes, and rumen face a similar biochemical challenge: the efficient degradation of complex organic matter under extreme physicochemical constraints. Each system represents a unique evolutionary solution to biomass conversion, shaped by distinct pressures yet converging on microbial enzymatic adaptations that enable organic matter breakdown. Both ecosystems host dense and metabolically active microbial consortia, specialized in enzymatic transformations related to carbon cycling, fiber degradation, and energy production^10–14^. These parallels underscore how distinct environmental pressures have driven convergent microbial strategies to solve a common metabolic challenge.

In soda lakes, microbial communities rely on dissolved inorganic carbon, primarily bicarbonate and carbonate, as the basis for primary production. Photosynthetic microbes, particularly cyanobacteria, convert these compounds into organic biomass, while heterotrophic alkaliphiles metabolize dissolved organic carbon derived from decaying microbial mats and extracellular polymers^4,15,16^. In contrast, the rumen microbiome specializes in the anaerobic fermentation of plant-derived polysaccharides, including cellulose, hemicellulose, lignin, and starch. These polymers are broken down into volatile fatty acids (VFAs) such as acetate, propionate, and butyrate, which serve as the primary energy source for the host animal. The rumen microbiome’s enzymatic repertoire includes fibrolytic enzymes adapted to low-oxygen conditions, functioning through syntrophic interactions among bacteria, protozoa, and methanogens ^14,17^.

Previous studies on the microbial composition and enzymatic potential of Ethiopia’s soda lakes have primarily relied on culture-dependent approaches, which capture only a limited subset of microbial diversity and functional capacity. More recently, culture-independent methods have significantly broadened our understanding of the taxonomic landscape of soda lake microbiomes^6,18^. For example, our recent amplicon-based metagenomic study revealed one of the highest recorded abundances of prokaryotic and eukaryotic taxa in soda lake ecosystems. Complementing this, a functional metagenomic analysis uncovered novel carbohydrate-active enzymes (CAZymes), highlighting the potential of these extreme environments as reservoirs of biotechnologically valuable biocatalysts^13,19,20^.

While these studies have advanced our understanding of microbial communities and their roles in carbohydrate degradation under extreme conditions, they also present notable limitations. Amplicon sequencing, although effective for taxonomic profiling, offers limited insight into functional potential and metabolic interactions^21,22^. Conversely, functional metagenomics excels at identifying specific enzymes but lacks the capacity to comprehensively characterize microbial community structure and ecosystem-level functionality^23–25^. To avoid these limitations, shotgun metagenomics enables comprehensive taxonomic and functional characterization, allowing for the simultaneous identification of microbial lineages, metabolic pathways, and enzymatic capabilities^23,26^.

Studying the microbial communities of both soda lakes and the rumen of ruminants presents a unique opportunity to uncover enzymes with broad environmental tolerance and novel catalytic mechanisms. Despite the growing availability of metagenomic datasets, comparative analyses that link enzyme function to environmental adaptation across distinct ecosystems remain limited. This study addresses that gap by integrating ecological context with functional potential through shotgun metagenomics. Such comparisons not only illuminate how selective pressures shape microbial enzyme repertoires but also enable the discovery of functionally analogous yet structurally novel enzymes. Ultimately, this work advances our understanding of microbial diversity and enzymatic innovation, with broad implications for ecological theory and biotechnological applications.

## Materials and Methods

### Sampling Sites and Collection Procedures

Water and sediment samples were collected from three East African Rift Valley soda lakes: Lakes Abijata, Chitu, and Shala, as described in Jeilu et al^20^. Water samples were obtained using sterile Niskin bottles, while sediment samples were collected using polyethylene bags. Water samples were filtered within 24 hours through polycarbonate membranes (22 μm pore size, 47 mm diameter; GE) to collect microbial biomass for DNA extraction. Rumen content samples were aseptically collected from freshly slaughtered goats, cattle, and sheep at the Addis Ababa Abattoirs Enterprise. Immediately post-mortem, approximately 50 mL of rumen content was extracted from each animal using sterile instruments and transferred into sterile 50 mL Falcon tubes. Samples were immediately placed on ice and transported to the laboratory under cold chain conditions. All samples were processed within two hours of collection to preserve microbial integrity. The pH of all samples was measured using a pH meter (OAKTON-pH110), and the salinity of soda lake samples was measured using a refractometer (HHTEC).

### DNA Extraction

DNA from lake water and sediment samples was extracted using the CTAB (cetyltrimethylammonium bromide)/SDS (sodium dodecyl sulfate) method, as described in our previous study ^20^. This protocol was optimized for high-yield and high-purity DNA recovery from environmental samples collected from soda lakes.

DNA extraction from rumen samples (goat, cattle, and sheep) was performed using the DNeasy PowerSoil Kit (QIAGEN), following the manufacturer’s instructions. Briefly, 0.25 g of each sample was transferred into a PowerBead Tube containing lysis buffer and subjected to bead beating using a vortex adapter. The supernatant was separated by centrifugation, and inhibitors were removed using Inhibitor Removal Technology (IRT) reagents. DNA was then bound to a silica membrane within a spin column, washed to remove contaminants, and eluted in 50 μL of TE buffer (pH 8.0). DNA quality and concentration were assessed using a NanoDrop spectrophotometer (Thermo Scientific) and agarose gel electrophoresis.

### Shotgun Metagenomic Sequencing

Shotgun metagenomic sequencing was performed by Novogene using the Illumina platform (PE150 reads, paired-end 150 bp). Libraries were prepared according to Novogene’s metagenomics library preparation protocol to ensure high-quality sequencing output. Each sample was sequenced to a minimum depth of 5 gigabases (Gb). Quality control assessments ensured that the proportion of bases with a Q30 score (≥85%) was maintained to guarantee high sequencing accuracy. Raw sequencing reads underwent standard quality filtering, adapter trimming, and preprocessing before downstream bioinformatics analyses, as described below.

### Quality Control and Contaminant Removal

Raw sequencing reads were assessed using FastQC^27^ to evaluate sequence quality. Adapters and low-quality bases were trimmed using Trimmomatic ^28^ with the parameters SLIDINGWINDOW:4:20 MINLEN:50. Contaminant sequences, including human DNA and PhiX control DNA, were filtered using KneadData (https://huttenhower.sph.harvard.edu/kneaddata/) under default parameters. Overrepresented sequences were identified and removed. A final quality check was conducted using FastQC, and summary reports were compiled using MultiQC ^29^. Sequencing coverage effort analysis was performed using Nonpareil v3.5.5^30^ to estimate microbial diversity and evaluate sequencing depth across samples.

### Read-Based Microbial Community Profiling and Functional Analysis

Taxonomic profiling of high-quality metagenomic reads was conducted using MetaPhlAn4 ^31^, employing the Jun23_CHOCOPhlAnSGB_202307 database. The resulting taxonomic profiles were analyzed to assess microbial community composition and diversity indices. Functional annotation of the metagenomic reads was performed using HUMAnN 3^32^, with gene families mapped to MetaCyc reactions using predefined HUMAnN mapping files. The data were normalized to relative abundance, and non-stratified tables were used for further analyses and visualizations.

### Metagenomic Assembly and Genome Construction

High-quality metagenomic reads were assembled into contigs using MEGAHIT v1.2.9^33^ with default parameters. Assembly quality was assessed using QUAST v5.2.0^34^. MAGs were reconstructed using MetaBAT 2^35^ by aligning reads to the assembled contigs using Bowtie2 v2.5.4^36^. BAM files were sorted and indexed using Samtools v1.20 ^37^. The completeness and contamination of the assembled MAGs were assessed using CheckM v1.1.6^38^, and only MAGs with ≥50% completeness and ≤10% contamination were retained for downstream analyses.

### Taxonomic and Functional Annotation of MAGs

Taxonomic classification of high-quality MAGs was performed using GTDB-Tk v2.4.0 ^39^, with the most recent GTDB Release 220. In addition to taxonomic assignment, Relative Evolutionary Divergence (RED) values were obtained from GTDB-Tk outputs and used to assess the evolutionary novelty of the reconstructed MAGs. Functional annotation was conducted using Prokka v1.14.5^40^ to identify coding sequences, rRNAs, and tRNAs. We used EcoFoldDB (v1.3), a structure-guided annotation tool that integrates Foldseek for structural alignment and ProstT5 for protein embedding, to functionally annotate predicted protein sequences from each MAG^41,42^. Carbohydrate-active enzymes (CAZymes)^43^ were identified using dbCAN2, which integrates HMMER, DIAMOND, and Hotpep for CAZyme classification. Codon profiling across MAGs, was performed using EMBOSS cusp ^44^ to calculate the frequency and relative fraction of each codon per amino acid, as well as GC content at the first, second, and third codon positions (GC1, GC2, GC3).

A phylogenetic tree of high-quality MAGs was constructed using PhyloPhlAn v3.0^45^ with default parameters. The tree was visualized using ggtree (Bioconductor R package).

## Statistical Analysis and Visualization

All downstream statistical analyses and visualizations were conducted in R v4.3.1^46^ using the vegan, phyloseq, ggplot2, ComplexHeatmap, and patchwork packages. Shannon diversity and Bray-Curtis dissimilarity were calculated to assess alpha and beta diversity, respectively. PCoA plots were used to visualize community and CAZyme compositional differences. Heatmaps and plots were used to display taxonomic, functional, and enzymatic abundance patterns. Genus-level contributions to glycoside hydrolase families were assessed using bar plots based on MAG annotations. The RSCU values were computed from codon frequencies using custom R scripts and averaged by sample type to identify environment-specific codon preferences. Statistical comparisons between groups were performed using the Kruskal-Wallis test, Wilcoxon rank-sum test, and PERMANOVA, with multiple-testing correction applied where appropriate.

## Results

### Microbial Composition of the Rumen and Soda Lakes

A total of 34 samples were analyzed, comprising 12 from soda lakes and 22 from rumen sources. A total of 1,007,699,292 clean sequence reads were obtained across all samples, with read counts varying across different sample types. The majority of samples approached high estimated average coverage, indicating sufficient sequencing depth to capture dominant community members (Supplementary Table 1, Supplementary Figure 1).

Proteobacteria and Cyanobacteria were more prominent in soda lake samples, while Firmicutes, Bacteroidota, and Fibrobacteres appeared to dominate rumen samples. Certain phyla such as Euryarchaeota and Actinobacteria also showed habitat-specific distributions, with notable presence in soda lakes. At the genus level, soda lakes were dominated by halotolerant and alkaliphilic genera such as *Halomonas*, *Nitrincola*, *Limnospira*, and multiple unclassified genera (e.g., GGB73217, GGB23825), with variable patterns across lake sites. In contrast, rumen samples were enriched in genera such as *Prevotella*, *Fibrobacter*, and GGB30811, consistent with microbial taxa involved in fermentative digestion of plant material. The presence of numerous unclassified genera across both environments further suggests the existence of novel or poorly characterized microbial taxa (Figure 1).

**Figure 1:**
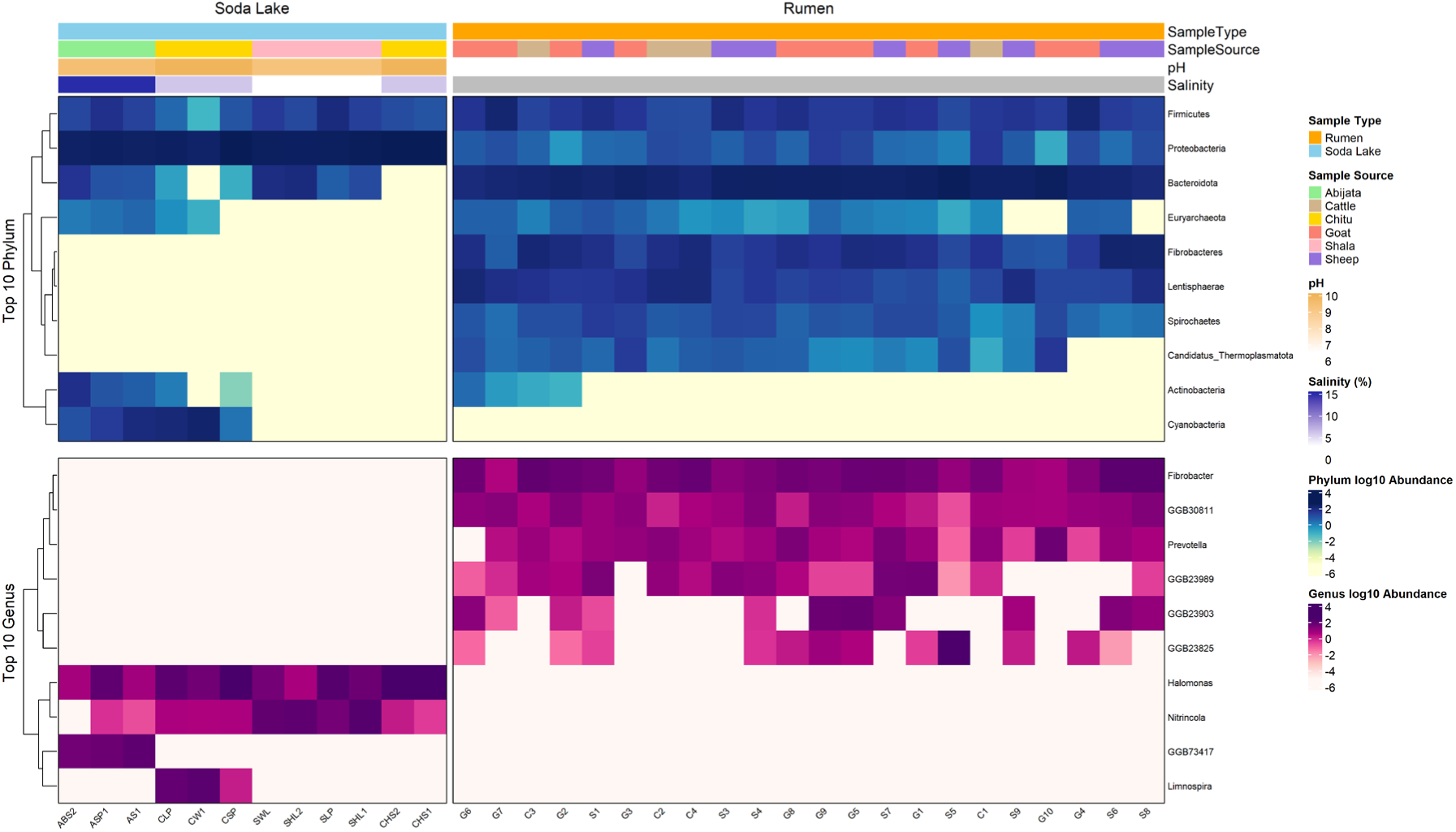
Heatmap showing the distribution of the top 10 most abundant microbial phyla (top panel) and genera (bottom panel) across soda lake and rumen samples. Samples are grouped by environment (Soda Lake vs Rumen), and annotated with metadata including sample source, pH, and salinity. Taxa abundance is shown as log₁₀-transformed values. Note: salinity was not measured for rumen samples, represented in gray.

### Microbial Diversity Indices of the Rumen and Soda Lakes

Microbial diversity differed significantly between soda lake and rumen environments. Rumen samples exhibited higher Shannon diversity compared to soda lake samples, with Chitu showing the lowest within the soda lake group (Figure 2, left panel). Among rumen sources, Sheep and Goat samples had relatively higher diversity than Cattle. Beta diversity analysis using PCoA based on Bray–Curtis distances revealed clear separation between soda lake and rumen samples (Figure 2, right panel). PERMANOVA analysis confirmed a statistically significant difference in community composition between the two environments (p = 0.001, R² = 0.17), highlighting distinct microbial assemblages shaped by host-associated and extreme lake conditions (Figure 2).

**Figure 2:**
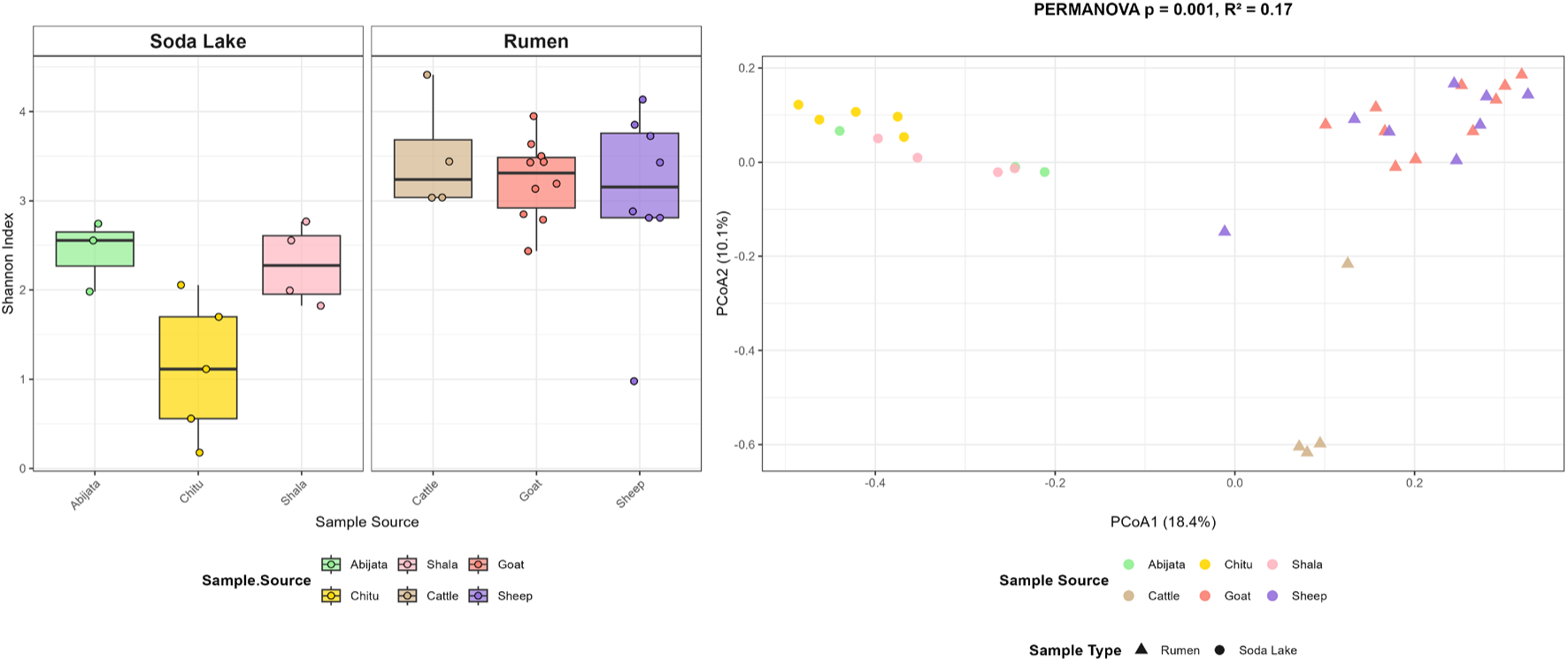
Alpha and beta diversity of microbial communities in soda lake and rumen samples. Left panel: Boxplot of Shannon diversity index grouped by sample source and environment (Soda Lake vs Rumen). Right panel: Principal coordinates analysis (PCoA) based on Bray–Curtis distances showing distinct clustering by sample type.

### Functional Potential of Microbial Communities

Pathways such as phosphoenolpyruvate carboxykinase (4.1.1.49), involved in gluconeogenesis, and pyruvate, phosphate dikinase (2.7.9.1), crucial for anaerobic respiration, were highly abundant in rumen samples. Additionally, amino acid biosynthesis pathways, including tryptophan synthase (4.2.1.20) and phenylalanine--tRNA ligase (6.1.1.20), were significantly enriched, supporting microbial protein synthesis and nitrogen assimilation. In addition, an abundance of endo-1,4-beta-xylanase (3.2.1.8), an enzyme essential for hemicellulose degradation, was observed. In contrast, the soda lake microbiome displayed functional adaptations suited for extreme environmental conditions, with strong enrichment in osmotic stress tolerance, sulfur metabolism, and genome maintenance pathways such as DNA helicase (3.6.4.12), DNA-directed DNA polymerase (2.7.7.7) and methionyl aminopeptidase (3.4.11.18) (Figure 3).

**Figure 3:**
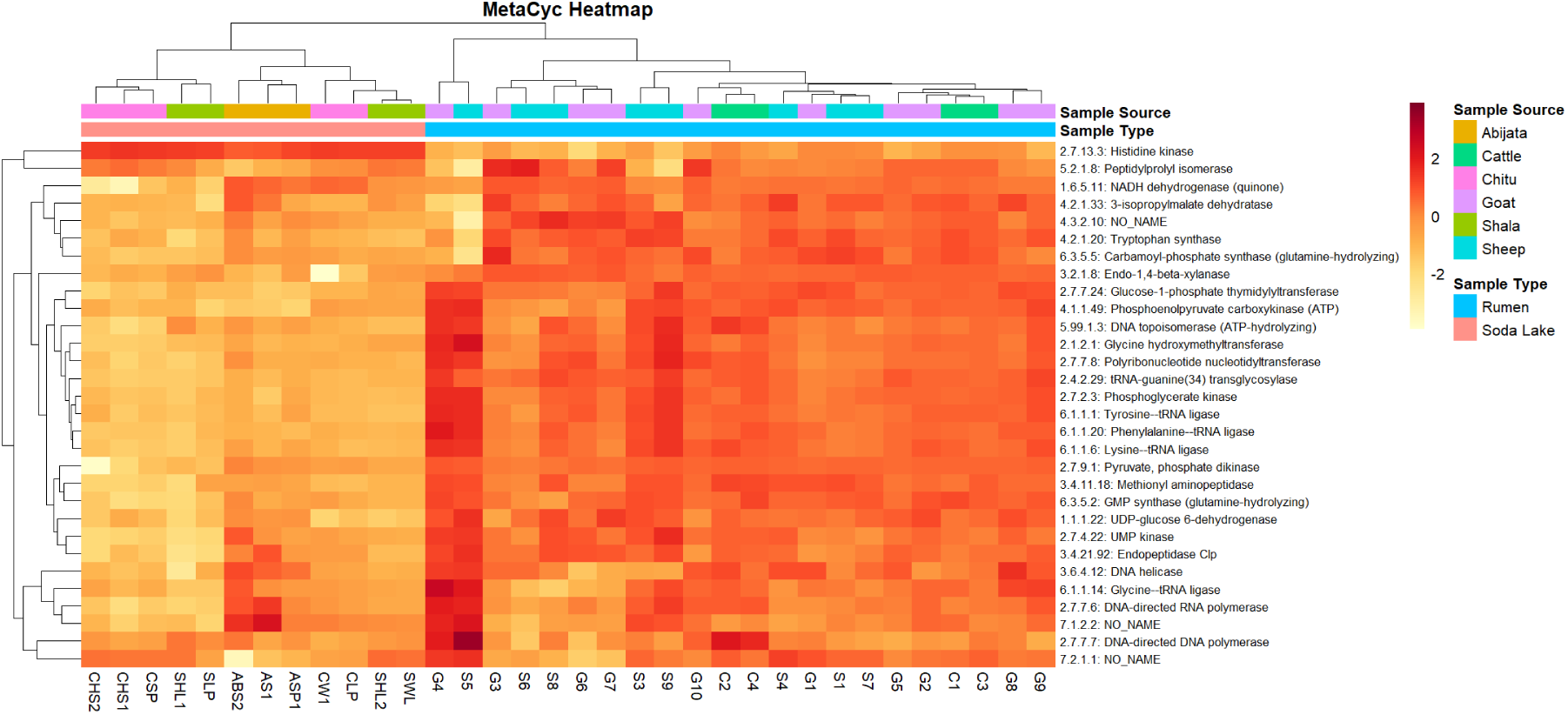
MetaCyc Pathway Heatmap: Hierarchical clustering heatmap displaying functional pathway enrichment across microbial communities from different sample sources. The color gradient represents pathway abundance, with red indicating high activity and yellow indicating lower activity.

### Genomic Characteristics and Evolutionary Insights of Metagenome-Assembled Genomes (MAGs)

A total of 1,208 metagenome-assembled genomes (MAGs) were reconstructed from the metagenomic datasets. MAGs were filtered based on ≥50% completeness and ≤10% contamination, resulting in 500 high-quality MAGs. These high-quality MAGs were distributed almost evenly across the two sample types, with 252 MAGs from rumen samples and 248 MAGs from soda lake samples. Most MAGs demonstrated high completeness (>90%) and low contamination (<5%), as visualized in the CheckM-based quality assessment (Figure 4A).

**Figure 4:**
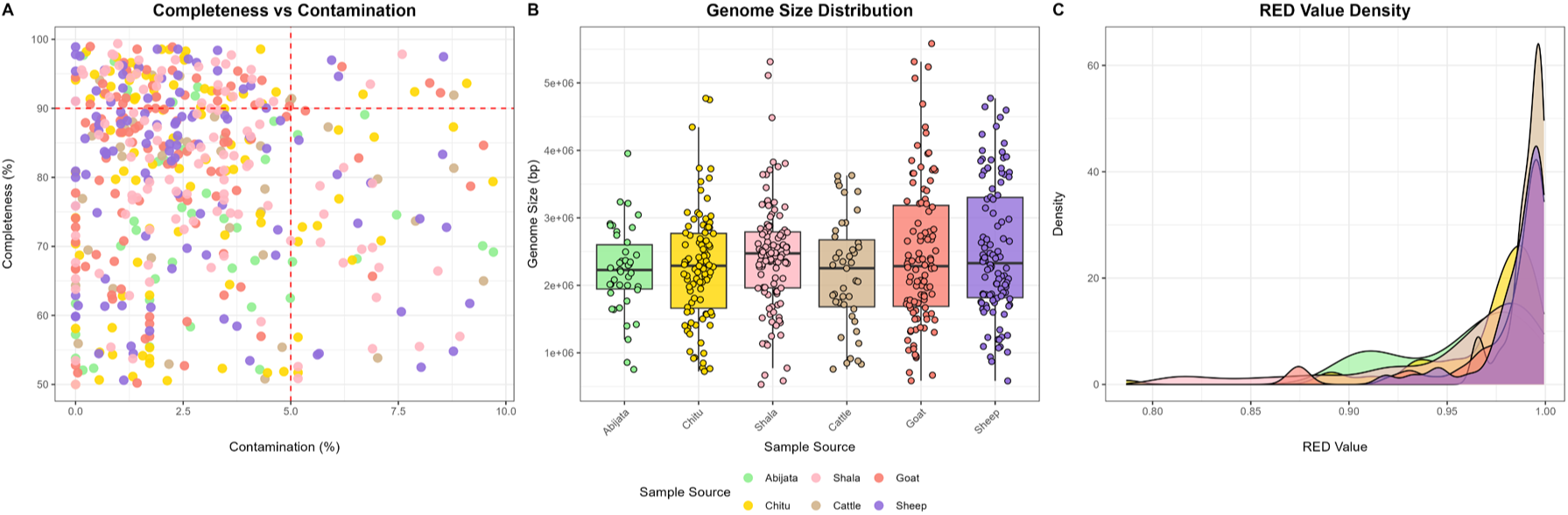
Quality, genome size, and RED value distribution of MAGs recovered from soda lake and rumen samples. (A) Scatterplot of MAG completeness vs. contamination (CheckM), colored by sample source. (B) Boxplot showing the distribution of estimated genome sizes (in base pairs) across sample sources. (C) Density plot of Relative Evolutionary Divergence (RED) values across sample sources

Genome size distributions varied among sample sources, with larger genomes more frequently recovered from Goat and Sheep rumen samples compared to the soda lake samples, particularly Chitu (Figure 4B). Relative Evolutionary Divergence (RED) value density plots indicated that the MAGs from soda lakes, especially Chitu and Abijata, harbored a higher proportion of evolutionarily divergent lineages (i.e., higher RED values) compared to rumen MAGs (Figure 4 C), suggesting the presence of phylogenetically novel taxa in these extreme environments.

### Taxonomic Composition of the Metagenome-Assembled Genomes (MAGs)

The phylogenetic relationships of the 500 high-quality MAGs reconstructed from rumen and soda lake environments were inferred using PhyloPhlAn, with taxonomic annotations based on GTDB-Tk classifications, among these 387 MAGS were successfully constructed in the tree (Figure 5).

**Figure 5:**
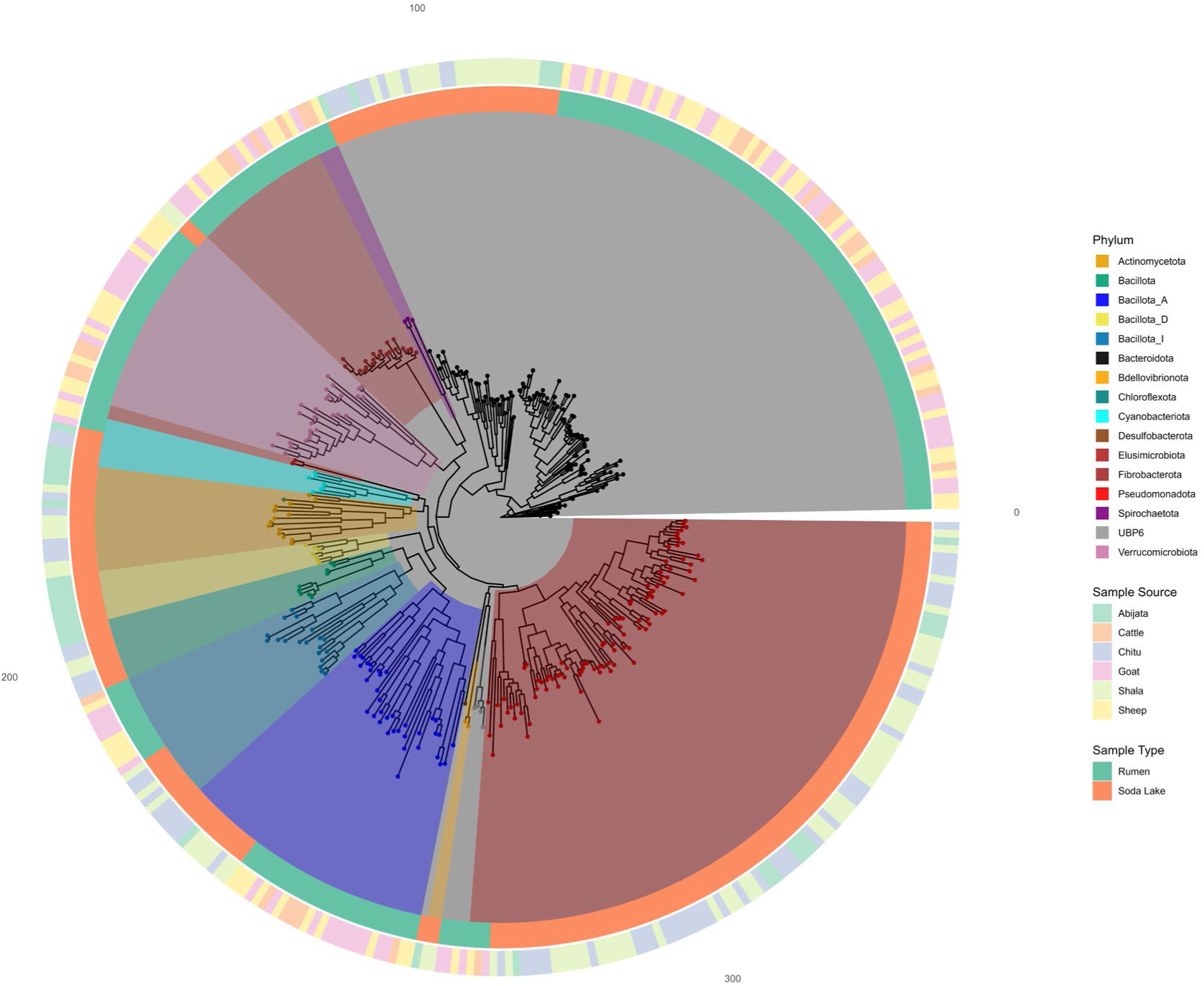
Phylogenetic tree of metagenome-assembled genomes (MAGs) reconstructed using PhyloPhlAn. Clades are color-coded based on phylum-level taxonomy according to GTDB-Tk classifications. Outer rings represent metadata for sample type (Rumen vs. Soda Lake) and sample source (Abijata, Chitu, Shala, Cattle, Goat, Sheep).

At the phylum level, MAGs from soda lake samples were dominated by Pseudomonadota and Bacteroidota, followed by Verrucomicrobiota, Fibrobacterota, and Bacillota_A, with several candidate and less-characterized phyla also represented (e.g Bdellovibrionota). In contrast, rumen-derived MAGs were enriched in Bacteroidota, Fibrobacterota, and Bacillota_I, along with Actinomycetota and Elusimicrobiota, consistent with the fibrolytic and fermentative roles typical of ruminant microbial communities. At the genus level, the rumen samples were largely characterized by well-known gut-associated genera, including *Fibrobacter*, *Prevotella*, and *Cryptobacteroides*, with the highest counts observed in sheep and goat samples. In contrast, soda lake samples harbored a broader array of genera, many of which were affiliated with extremophiles or uncultured taxa, including *Aliidiomarina*, *Marinospirillum*, *Natronospirillum*, and *Alkalibacterium*. Notably, a substantial fraction of MAGs remained unclassified at the genus level, particularly in soda lake datasets, suggesting the presence of potentially novel or under-characterized microbial lineages (Figure 6).

**Figure 6:**
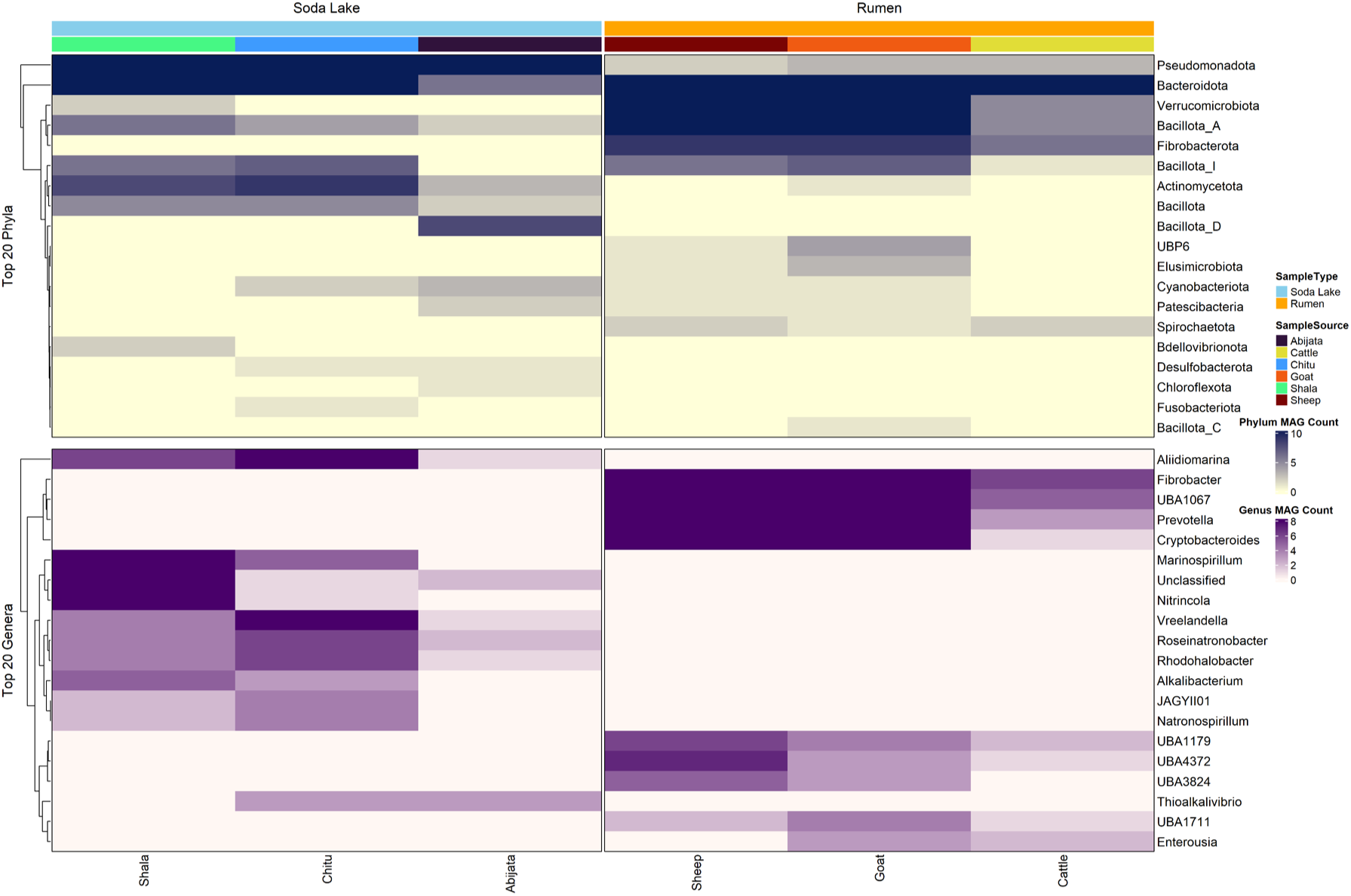
Heatmap showing the top 20 most abundant phyla (top panel) and genera (bottom panel) assigned to metagenome-assembled genomes (MAGs) recovered from Soda Lake (left) and Rumen (right) environments. MAG abundance is represented by color intensity, with hierarchical clustering highlighting distinct taxonomic profiles across sample sources.

### Functional potential of the Metagenome-Assembled Genomes (MAGs)

Carbon cycling dominated the functional category composition across all samples, comprising over 60% of annotated functions, followed by sulfur cycling, osmotic stress tolerance, and nitrogen cycling. The relative abundance of non-carbon functional categories was slightly higher in soda lake samples, particularly for sulfur cycling and osmotic stress tolerance, reflecting adaptation to high-salinity and alkaline environments (Figure 7A). In the Carbon cycle, Rumen samples were enriched in hemicellulose, cellulose, and starch degradation, whereas soda lakes showed higher relative abundances of aromatic compound metabolism pathways such as toluene conversion, ferulic acid to vanillin conversion, and caffeate respiration. These differences suggest divergent strategies for organic matter turnover between host-associated and extremophilic microbial communities (Figure 7B). Glycoside hydrolases (GH) and glycosyltransferases (GT) dominated all samples, but auxiliary activities (AA) and carbohydrate-binding modules (CBM) were relatively more abundant in soda lake samples. Glycosyltransferases (GTs) were the most abundant CAZyme class in all samples. Glycoside hydrolases (GHs) ranked second and were particularly enriched in rumen samples (goat and sheep), highlighting their role in complex carbohydrate degradation in host-associated microbiomes (Figure 7C).

**Figure 7:**
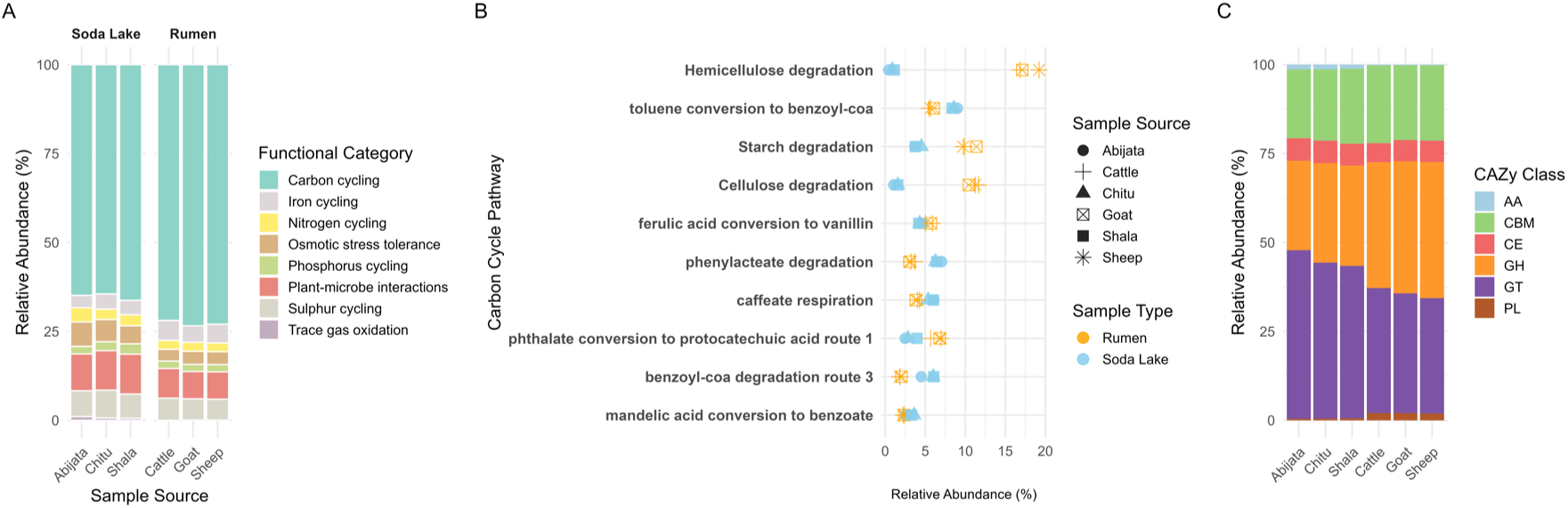
Functional category, pathway-level, and enzymatic class comparisons between soda lake and rumen microbiomes. (A) Relative abundance of functional categories derived from MAG annotations, grouped by sample type. (B) Dot plot showing the relative abundance of the top 10 carbon cycle pathways across all samples. The dot color denotes the sample type; the dot shape denotes the individual sample source. (C) CAZy class composition based on annotated CAZyme families from metagenomic assemblies. Samples are ordered consistently across panels.

**Figure 8:**
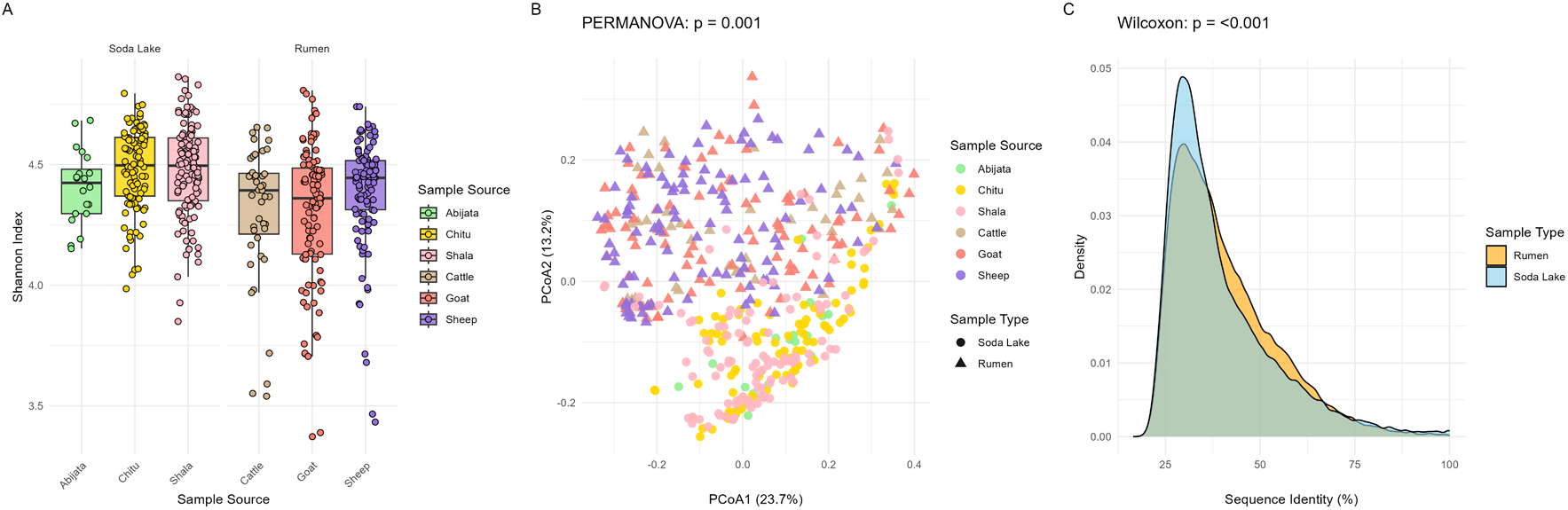
Functional profiling of CAZymes across soda lake and rumen metagenomes. (A) Shannon diversity indices indicate slightly higher CAZyme diversity in soda lake samples compared to rumen. (B) Principal Coordinates Analysis (PCoA) based on Bray-Curtis dissimilarity reveals distinct clustering of CAZyme profiles by the environment. (C) Density plot of sequence identity (%) between predicted CAZymes and CAZy reference entries, showing a higher proportion of divergent CAZymes in soda lake samples.

The rumen MAGs exhibited significantly higher GC content at the third codon position (GC3) compared to those from soda lakes (Wilcoxon rank-sum test, p = 1e−5). To assess codon usage bias independent of amino acid composition, we calculated Relative Synonymous Codon Usage (RSCU) values across MAGs from both environments. Codons ending in G or C were more frequently used in rumen MAGs, consistent with their elevated GC3 content and stronger codon bias. In contrast, soda lake MAGs preferentially used A- or T-ending codons for several amino acids, reflecting a distinct codon usage pattern potentially shaped by mutational bias or relaxed translational selection (Supplementary Fig. 2).

### CAZy Gene Family Diversity and Functional Richness

At the diversity level, soda lake samples exhibited moderately higher CAZyme richness and evenness compared to rumen samples, as indicated by the Shannon Index (Figure 7A). The highest CAZyme diversity was observed in Chitu and Shala lakes, while Abijata showed slightly lower values. Among rumen sources, sheep and goat samples harbored more diverse CAZyme repertoires than cattle, suggesting variability in carbohydrate degradation potential related to host species or diet. Principal Coordinates Analysis (PCoA) clearly separated soda lake and rumen samples, with minimal overlap, indicating strong functional divergence in CAZyme composition. Within each environment, source-specific grouping further highlighted localized ecological structuring of functional gene content (Figure 7B). Notably, soda lake samples contained a broader range of low-identity CAZymes, reflecting a higher degree of novelty or divergence. This pattern suggests that soda lake microbiomes may encode unique or previously uncharacterized carbohydrate-active enzymes, potentially adapted to extreme haloalkaline conditions (Figure 7C).

Among the GH families, GH13 (α-amylase, pullulanase) was the most abundant, especially in goat and sheep rumen samples, suggesting a strong capacity for starch degradation and α-glucan hydrolysis in host-associated communities. This was followed by GH2 (β- galactosidase, mannosidase) and GH3 (β-glucosidase, arabinofuranosidase), both broadly distributed but enriched in rumen samples, indicating an elevated ability to hydrolyze β- linked oligosaccharides and hemicellulose-derived sugars. GH43 (arabinofuranosidase, xylosidase) was also abundant in both environments, with a strong representation in Shala (soda lake) and goat/sheep (rumen), pointing to shared functional roles in xylan degradation. Other notable families include GH23 (peptidoglycan hydrolase), GH5 (cellulase, xylanase, mannanase), and GH1 (β-glucosidase), which were moderately represented in both environments. Lower abundance families, such as GH171, GH109, GH120, GH28, and GH97, were more variable but still contributed to the functional repertoire, with GH109 (N-acetylgalactosaminidase) being more prominent in soda lake samples (Figure 9).

**Figure 9:**
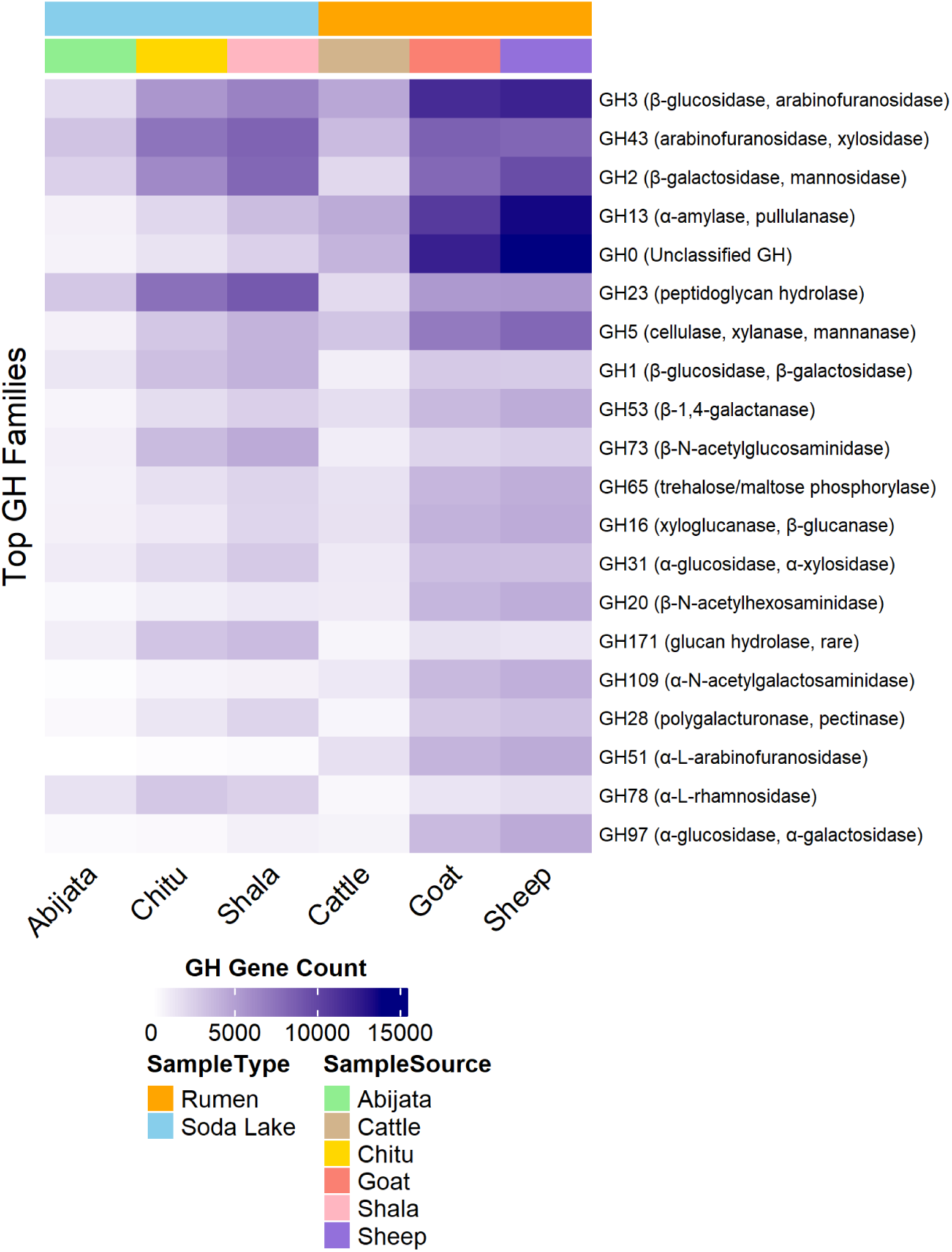
Heatmap of the top 20 most abundant glycoside hydrolase (GH) families across Soda Lake (Abijata, Chitu, Shala) and rumen (Cattle, Goat, Sheep) metagenomes. Gene abundance is indicated by color intensity, with darker shades representing higher counts.

**Figure 10:**
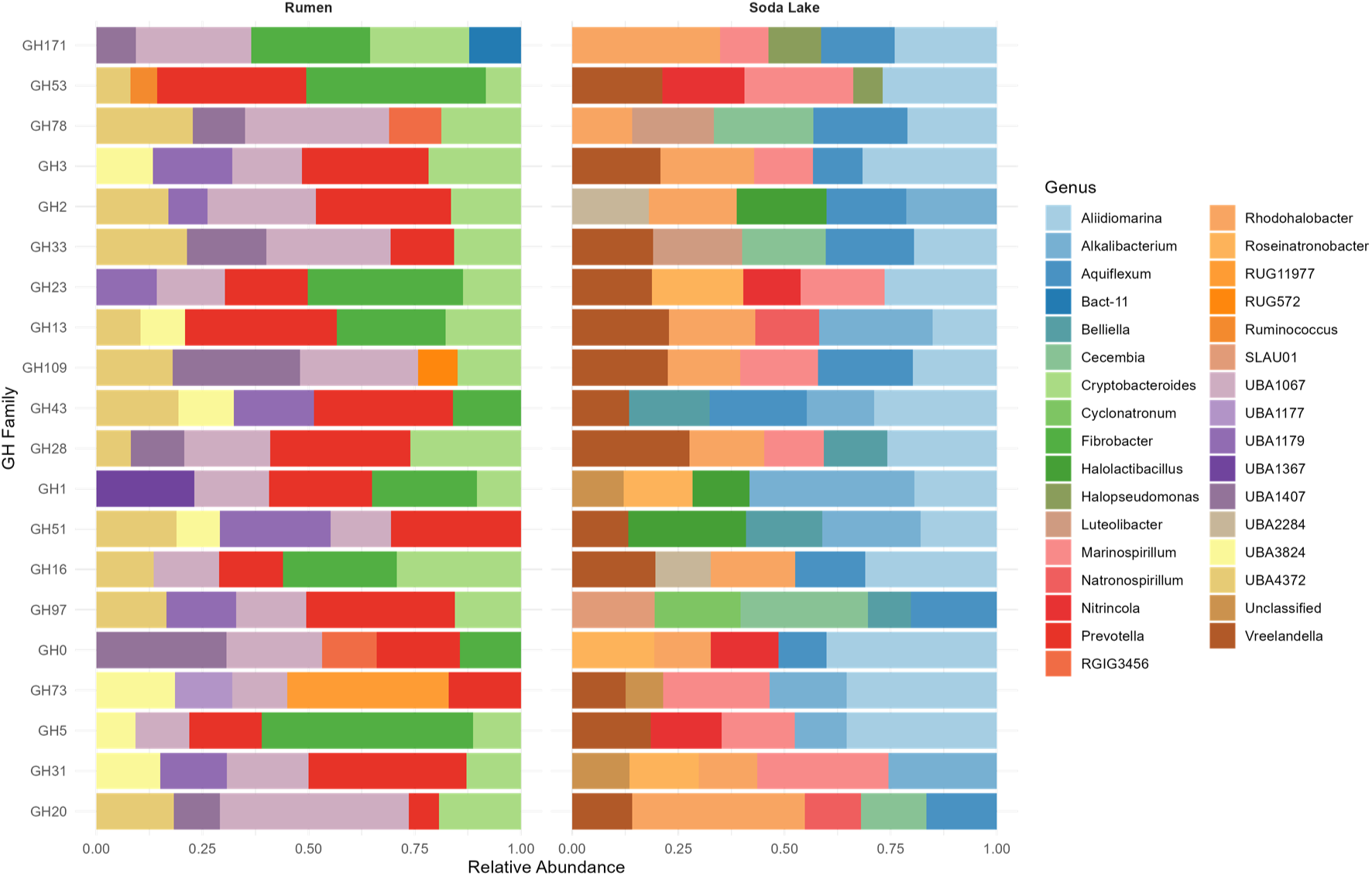
Relative abundance of the top 5 genera contributing to glycoside hydrolase (GH) families across two sample types: Rumen and Soda Lake. Each bar represents the composition of genera associated with a GH family, with proportions calculated based on relative abundance within each group. Taxonomic contributions were determined from genome-resolved annotations, and only the top 5 genera per GH family per sample type were included.

To identify microbial contributors to carbohydrate-active enzyme (CAZyme) activity across distinct environments, we examined the top five most abundant genera for selected glycoside hydrolase (GH) families associated with biomass degradation in rumen and soda lake samples. In the rumen microbiome, *Fibrobacter* and *Prevotella* were consistently dominant across multiple GH families. Other notable contributors included UBA1067, *Cryptobacteroides*, UBA1179, and *Ruminococcus*. In contrast, the soda lake microbiome showed a distinct set of dominant genera. *Alkalibacterium* was highly abundant in many of GH families. *Aliidiomarina* was among the top genera for GH1, GH3, GH5, and GH43, suggesting its broad role in polysaccharide degradation. Additional key genera included *Rhodohalobacter*, *Aquiflexum*, *Salisediminibacterium*, and *Isachenkonia*.

## Discussion

Understanding microbial diversity and functional adaptations in extreme environments is essential for uncovering novel metabolic pathways and biotechnological resources^47^. Here, we present, to our knowledge, the first comprehensive profile of carbohydrate-degrading enzymes present in microbial communities from two ecologically and functionally contrasting environments: the ruminant digestive tract and Ethiopian soda lakes. While the rumen microbiome, well-studied for its role in fiber degradation and fermentation^9,48^, is characterized by mildly acidic, low salt conditions, soda lakes are characterized by high alkalinity and salinity^15,49^, and their microbial diversity remains largely unexplored.

In our study, the microbial communities in the rumen and soda lakes exhibited distinct taxonomic profiles and diversity patterns which are likely shaped by their contrasting ecological roles and environmental conditions. Rumen samples displayed significantly higher microbial diversity (median Shannon index ∼4.5) compared to soda lake samples (∼3.2; p < 0.001), consistent with the rumen’s function as a complex, cooperative digestive system where microbial consortia work synergistically to degrade plant material^50,51^. Beta diversity analysis using Principal Coordinates Analysis (PCoA) based on Bray-Curtis dissimilarity revealed a strong separation between rumen and soda lake microbiomes (PERMANOVA p = 0.001), confirming that microbial composition is primarily driven by environmental selection^52,53^.

The reconstruction of 1,208 (500 high-quality, with 252 from rumen and 248 from soda lakes) MAGs from our metagenomic dataset offers unprecedented insights into microbial diversity, genome architecture, and evolutionary strategies in these two contrasting ecosystems. The observed differences in genome size distributions between rumen and soda lake microbiota in this study highlight how environmental pressures shape microbial genomic architecture. Rumen MAGs display a wide range of genome sizes, from approximately 1 Mbp to over 6 Mbp, similar to other studies^54–56^. This diversity reflects the complex metabolic capabilities required to degrade various plant polysaccharides and facilitate host-associated processes. In contrast, soda lake MAGs exhibit a narrower genome size distribution, typically between 2 to 4 Mbp. This suggests selective pressures favoring metabolic efficiency and specialized adaptations to high-pH and high-salinity conditions. This phenomenon aligns with the genome streamlining observed in extremophiles, where genomes are reduced to eliminate redundant functions while retaining essential genes for survival in harsh environments^57,58^.

The observed differences in GC3 content and codon usage bias between rumen and soda lake MAGs reflect distinct evolutionary adaptations to their respective environments. Rumen microbes, residing in a stable, nutrient-rich, and anaerobic host environment, exhibit higher GC3 content and a preference for G/C-ending codons. This likely enhances translational efficiency and genomic stability, aligning with the host’s metabolic demands and the availability of specific tRNAs ^59^. In contrast, soda lake microbes, exposed to high salinity, alkalinity, and UV radiation, show lower GC3 content and favor A/T-ending codons. This pattern may result from mutational biases driven by environmental stress and the energetic advantage of synthesizing A/T nucleotides ^60^.

This study’s microbial composition and comparative analysis of MAGs from rumen and soda lake environments reveal distinct taxonomic compositions, reflecting the unique selective pressures shaping microbial communities in each ecosystem. The rumen microbiome is dominated by *Bacteroidaceae*, and *Fibrobacteraceae*. These families are known for their ability to break down complex carbohydrates derived from plant material, contributing to host nutrition and energy production through fermentation processes^54,61^. *Prevotella*, *Fibrobacter*, *Cryptobacteroides,* and *Ruminococcus* were among the most abundant genera in the rumen, reflecting their established role in carbohydrate fermentation and lignocellulose degradation^62,63^. *Fibrobacter* being a primary cellulose degrader, and *Prevotella* contributes significantly to hemicellulose breakdown and overall carbohydrate metabolism^64–67^. *Cryptobacteroides* also contribute to rumen function, specifically carbohydrate degradation, and are more abundant in cattle with lower carcass traits^68,69^.

In stark contrast, the soda lake microbiome exhibited a strong enrichment of *Halomonadaceae*, *Alteromonadaceae*, *Rhodobacteraceae*, and *Balneolaceae* families known for their adaptations to hypersaline and alkaline conditions^70–72^. *Halomonas*, *Nitrincola*, and *Limnospira* were enriched in soda lakes, consistent with their known adaptation to hypersaline and alkaline conditions^4,49^. In addition *Thioalkalivibrio*, a genus associated with sulfur oxidation, was detected in soda lake samples, underscoring the role of sulfur cycling in these extreme environments^73,74^.

In our study, the substantial proportion of unclassified microbial reads and the broader RED distribution observed in soda lake MAGs underscore the presence of novel, evolutionarily distinct microbial lineages in these environments. This finding aligns with studies indicating that soda lakes harbor diverse microbial communities with a significant fraction of unclassified taxa^75,76^. In our recent study on amplicon metagenomic study of these soda lakes, a notable portion remains unclassified at the species level ^49^, highlighting the potential for discovering novel microbial taxa in these extreme environments. In contrast, the rumen microbiome exhibits a more constrained RED range, reflecting a higher degree of relatedness to known microbial taxa, consistent with its well-characterized nature and functional specialization in plant polysaccharide degradation^55,77^. This pattern suggests that soda lakes serve as evolutionary hotspots for microbial novelty while the rumen microbiome maintains functional stability through conserved microbial lineages.

This study also showed that the functional potential of the rumen and soda lake microbiomes reflects their distinct ecological roles and environmental constraints, with metabolic pathways enriched in each environment and aligning with their respective adaptations. The rumen microbiome showed strong enrichment in phosphoenolpyruvate carboxykinase (4.1.1.49) and pyruvate, phosphate dikinase (2.7.9.1), enzymes crucial for gluconeogenesis and anaerobic respiration ^78–80^, key metabolic strategies in fermentative digestion. The presence of endo-1,4-beta-xylanase (3.2.1.8), an enzyme essential for hemicellulose degradation, further underscores the role of rumen microbes in breaking down plant fibers into fermentable sugars, which are subsequently converted into volatile fatty acids to fuel host metabolism^81^.

In contrast, Soda Lake microbial communities exhibit distinct functional adaptations to survive in extreme alkaline and hypersaline environments. One of the most striking observations in soda lake samples is the enrichment of genome maintenance pathways, including DNA helicase (3.6.4.12), DNA-directed DNA polymerase (2.7.7.7), and methionyl aminopeptidase (3.4.11.18), which suggests strong selective pressure to maintain genome integrity against oxidative and environmental stressors common in hypersaline environments^82,83^. Additionally, the strong presence of sulfur metabolism pathways underscores their role in oxidizing reduced sulfur compounds to sustain microbial primary production in organic carbon-limited environments^84^.

Soda lake samples exhibited slightly higher CAZy diversity (median Shannon index ∼4.5) than rumen samples (∼4.3), which is surprising given that ruminants rely on highly specialized carbohydrate metabolism to break down complex plant polysaccharides. The rumen is well known for its dense microbial consortia that facilitate fiber degradation through an extensive repertoire of carbohydrate-active enzymes ^81,85^. However, our results suggest that soda lake microbiomes harbor a more functionally diverse CAZy gene pool.

The substantial proportion of low-identity CAZy genes (<40%) identified in soda lake samples underscores the potential for novel CAZymes in these extreme environments. This observation aligns with our previous functional metagenomic study, in which most identified CAZymes were novel^13^. These findings support that soda lakes can serve as hotspots for enzymatic innovation, likely driven by selective pressures such as high alkalinity, salinity, and nutrient limitations. The rumen is nevertheless an established environment for CAZyme discovery due to its specialization in complex plant polysaccharide degradation and host-associated microbial metabolism^86–88^. This highlights the potential for uncovering novel biocatalysts with industrial and biotechnological applications from both ecosystems, expanding the search for robust and efficient enzymes beyond host-associated microbiomes to extremes.

The rumen microbiome, known for its ability to degrade plant-derived polysaccharides, was enriched in GH families such as GH2 (β-galactosidases and mannosidases), GH3 (β- glucosidases and xylosidases), GH5 (endoglucanases and mannanases), GH9 (endoglucanases), and GH43 (arabinanases and xylosidases). These GH families, predominantly linked to *Ruminococcus*, *Prevotella*, *Fibrobacter*, and *Butyrivibrio*, underscore the crucial role of these microbial genera in fiber degradation and fermentation within ruminant digestive systems^89–91^. GH13 (α-amylases), GH23 (lysozymes), GH77 (4-α-glucanotransferases), and GH109 (α-N-acetylgalactosaminidases) were among the abundant GH in the soda lakes, which suggests adaptations for utilizing diverse carbohydrate sources, potentially linked to microbial strategies for energy acquisition in oligotrophic environments. Genera such as *Halomonas*, *Alkalilimnicola*, and *Nitrosomonas* were enriched in GH families adapted to osmotic and oxidative stress, supporting previous findings that soda lake microbes rely on unique carbohydrate degradation pathways optimized for high alkalinity and salinity^4,13,92,93^. In addition to these well characterized genera, a notable proportion of GHs were affiliated with uncultivated lineages, particularly those classified under UBA (Uncultured Bacterial and Archaeal) clades. All of these UBA- associated genera remain uncultivated^94^, highlighting the hidden enzymatic potential within these poorly understood groups.

The genomic and enzymatic diversity uncovered in this study represents a valuable resource for industrial enzyme discovery. Rumen and soda lake microbiomes encode CAZymes with complementary advantages: the former is well-established in lignocellulose degradation, while the latter harbors enzymes adapted to high pH, salinity, and oxidative stress. These traits are increasingly relevant to sectors such as bioenergy, wastewater treatment, and biomass valorization. Integrating metagenomics with biochemical validation offers a powerful strategy to accelerate the deployment of environmental enzymes in industrial applications. For example, a β-glucosidase previously characterized from this soda lake metagenome exhibited both halotolerance and glucose tolerance features advantageous for saccharification and biomass processing^19^. Collectively, these findings highlight the potential of metagenomic mining to uncover extremophile-adapted enzymes with high translational value for biotechnology.

## Conclusion

This study provides novel insights into the microbial diversity, genomic features, and functional adaptations of microbial communities from two ecologically distinct environments: the ruminant gut and Ethiopian soda lakes. While the rumen microbiome is known for its high diversity and specialization in plant polysaccharide degradation, soda lake communities exhibited distinct taxonomic and metabolic signatures shaped by extreme alkalinity and salinity. Notably, the slightly higher CAZy gene diversity observed in soda lakes, along with a substantial fraction of low-identity CAZymes (<40%), suggests the presence of highly divergent or novel enzymes uniquely adapted to extreme conditions. These findings are consistent with our previous discovery of novel glycoside hydrolases and polysaccharide lyases from these environments. Moreover, the broader RED score distribution among soda lake MAGs supports their role as reservoirs of microbial and enzymatic novelty. Together, these results underscore the biotechnological potential of both ecosystems and highlight the importance of continued functional exploration of their microbial gene pools.

## Acknowledgements

We thank the Ethiopian Wildlife Conservation Authority for granting access to sample from Lakes Abijata, Chitu, and Shala. We are also grateful to the Ethiopian Biodiversity Institute and the National Soil Testing Center, Ministry of Agriculture of Ethiopia, for permitting the transfer of genetic material. We acknowledge the staff at the Addis Ababa Abattoirs Enterprise for their support during rumen content collection. Computational resources were provided by UPPMAX (Uppsala Multidisciplinary Center for Advanced Computational Science) and Dardel at the PDC Center for High Performance Computing (KTH Royal Institute of Technology, Sweden).

## Author Contributions

O.J., E.H., E.A., E.J., and A.S. conceived and designed the study. O.J. performed sample collection, laboratory work, and computational analyses, and drafted the manuscript. All authors contributed to data interpretation, provided critical feedback, and approved the final version of the manuscript.

## Data Availability

All raw sequencing data have been deposited in the NCBI Sequence Read Archive (SRA) under BioProject accession number PRJNA1273195. Additional data supporting the findings of this study are available from the corresponding author upon reasonable request.

## Code Availability

Custom scripts used for data preprocessing, taxonomy and functional annotation, statistical analysis, and visualization are available at: https://github.com/OliyadJe/Soda-Lakes-and-Rumen-Metagenomics.

## References

1. Rekadwad, B. N. et al. Extremophiles: the species that evolve and survive under hostile conditions. 3 Biotech 13, 316 (2023).

2. Sayed, A. m., et al. Extreme environments: microbiology leading to specialized metabolites. Journal of Applied Microbiology 128, 630–657 (2020).

3. Sorokin, D. Y., Banciu, H. L. & Muyzer, G. Functional microbiology of soda lakes. Current Opinion in Microbiology 25, 88–96 (2015).

4. Sorokin, D. Y. et al. Microbial diversity and biogeochemical cycling in soda lakes. Extremophiles 18, 791–809 (2014).

5. Agembe, S., Ojwang, W., Olilo, C., Omondi, R. & Ongore, C. Soda Lakes of the Rift Valley (Kenya). in The Wetland Book: II: Distribution, Description and Conservation (eds. Finlayson, C. M., Milton, G. R., Prentice, R. C. & Davidson, N. C.) 1–11 (Springer Netherlands, Dordrecht, 2017). doi:10.1007/978-94-007-6173-5_150-2.

6. Simachew, A., Lanzén, A., Gessesse, A. & Øvreås, L. Prokaryotic Community Diversity Along an Increasing Salt Gradient in a Soda Ash Concentration Pond. Microb Ecol 71, 326–338 (2016).

7. Cammack, K. M., Austin, K. J., Lamberson, W. R., Conant, G. C. & Cunningham, H. C. RUMINNAT NUTRITION SYMPOSIUM: Tiny but mighty: the role of the rumen microbes in livestock production1. J Anim Sci 96, 752–770 (2018).

8. Matthews, C. et al. The rumen microbiome: a crucial consideration when optimising milk and meat production and nitrogen utilisation efficiency. Gut Microbes 10, 115– 132 (2018).

9. Cammack, K. M., Austin, K. J., Lamberson, W. R., Conant, G. C. & Cunningham, H. C. RUMINNAT NUTRITION SYMPOSIUM: Tiny but mighty: the role of the rumen microbes in livestock production1. J Anim Sci 96, 752–770 (2018).

10. Barrett, K. et al. Changes in the Metagenome-Encoded CAZymes of the Rumen Microbiome Are Linked to Feed-Induced Reductions in Methane Emission From Holstein Cows. Front. Microbiol. 13, (2022).

11. Hinsu, A. T. et al. Characterizing rumen microbiota and CAZyme profile of Indian dromedary camel (Camelus dromedarius) in response to different roughages. Sci Rep 11, 9400 (2021).

12. Auer, E. et al. Horizontal metaproteomics and CAZymes analysis of lignocellulolytic microbial consortia selectively enriched from cow rumen and termite gut. ISME COMMUN. 3, 1–12 (2024).

13. Jeilu, O., Simachew, A., Alexandersson, E., Johansson, E. & Gessesse, A. Discovery of novel carbohydrate degrading enzymes from soda lakes through functional metagenomics. Front. Microbiol. 13, (2022).

14. Liang, J. et al. Rumen microbes, enzymes, metabolisms, and application in lignocellulosic waste conversion - A comprehensive review. Biotechnology Advances 71, 108308 (2024).

15. Zorz, J. K. et al. A shared core microbiome in soda lakes separated by large distances. Nat Commun 10, 4230 (2019).

16. Pellegrinetti, T. A. et al. The role of microbial communities in biogeochemical cycles and greenhouse gas emissions within tropical soda lakes. Science of The Total Environment 947, 174646 (2024).

17. Gheibipour, M. et al. Bioengineering the Rumen Microbiota as an Advanced Biocatalyst for Renewable Fuels. in Recent Trends in Lignocellulosic Biofuels and Bioenergy: Advancements and Sustainability Assessment (eds. Dar, M. A., Zabed, H. M. & Shahnawaz, M.) 143–175 (Springer Nature Singapore, Singapore, 2025). doi:10.1007/978-981-96-4636-4_6.

18. Lanzén, A. et al. Surprising Prokaryotic and Eukaryotic Diversity, Community Structure and Biogeography of Ethiopian Soda Lakes. PLOS ONE 8, e72577 (2013).

19. Jeilu, O., Alexandersson, E., Johansson, E., Simachew, A. & Gessesse, A. A novel GH3-β-glucosidase from soda lake metagenomic libraries with desirable properties for biomass degradation. Sci Rep 14, 10012 (2024).

20. Jeilu, O., Gessesse, A., Simachew, A., Johansson, E. & Alexandersson, E. Prokaryotic and eukaryotic microbial diversity from three soda lakes in the East African Rift Valley determined by amplicon sequencing. Frontiers in Microbiology 13, 1–14 (2022).

21. Alteio, L. V. et al. A critical perspective on interpreting amplicon sequencing data in soil ecological research. Soil Biology and Biochemistry 160, 108357 (2021).

22. Yi, X. et al. Unravelling the enigma of the human microbiome: Evolution and selection of sequencing technologies. Microb Biotechnol 17, e14364 (2023).

23. Robinson, S. L., Piel, J. & Sunagawa, S. A roadmap for metagenomic enzyme discovery. Nat Prod Rep 38, 1994–2023.

24. Mathieu, E. et al. An Insight into Functional Metagenomics: A High-Throughput Approach to Decipher Food–Microbiota–Host Interactions in the Human Gut. Int J Mol Sci 24, 17630 (2023).

25. Zuñiga, C., Zaramela, L. & Zengler, K. Elucidation of complexity and prediction of interactions in microbial communities. Microb Biotechnol 10, 1500–1522 (2017).

26. Usyk, M. et al. Comprehensive evaluation of shotgun metagenomics, amplicon sequencing, and harmonization of these platforms for epidemiological studies. Cell Reports Methods 3, 100391 (2023).

27. Andrews, S. FastQC. Babraham Bioinformatics (2010).

28. Bolger, A. M., Lohse, M. & Usadel, B. Trimmomatic: A flexible trimmer for Illumina sequence data. Bioinformatics 30, (2014).

29. Ewels, P., Magnusson, M., Lundin, S. & Käller, M. MultiQC: Summarize analysis results for multiple tools and samples in a single report. Bioinformatics 32, (2016).

30. Rodriguez-R, L. M. & Konstantinidis, K. T. Nonpareil: a redundancy-based approach to assess the level of coverage in metagenomic datasets. Bioinformatics 30, 629–635 (2014).

31. Blanco-Míguez, A. et al. Extending and improving metagenomic taxonomic profiling with uncharacterized species using MetaPhlAn 4. Nature Biotechnology 41, (2023).

32. Beghini, F. et al. Integrating taxonomic, functional, and strain-level profiling of diverse microbial communities with biobakery 3. eLife 10, (2021).

33. Li, D., Liu, C. M., Luo, R., Sadakane, K. & Lam, T. W. MEGAHIT: An ultra-fast single-node solution for large and complex metagenomics assembly via succinct de Bruijn graph. Bioinformatics 31, (2015).

34. Gurevich, A., Saveliev, V., Vyahhi, N. & Tesler, G. QUAST: Quality assessment tool for genome assemblies. Bioinformatics 29, (2013).

35. Kang, D. D. et al. MetaBAT 2: An adaptive binning algorithm for robust and efficient genome reconstruction from metagenome assemblies. PeerJ 2019, (2019).

36. Langmead, B. & Salzberg, S. L. Fast gapped-read alignment with Bowtie 2. Nature Methods. Nature Publishing Group (2012).

37. Danecek, P. et al. Twelve years of SAMtools and BCFtools. GigaScience 10, (2021).

38. Parks, D. H., Imelfort, M., Skennerton, C. T., Hugenholtz, P. & Tyson, G. W. CheckM: Assessing the quality of microbial genomes recovered from isolates, single cells, and metagenomes. Genome Research 25, (2015).

39. Chaumeil, P. A., Mussig, A. J., Hugenholtz, P. & Parks, D. H. GTDB-Tk: A toolkit to classify genomes with the genome taxonomy database. Bioinformatics 36, (2020).

40. Seemann, T. Prokka: Rapid prokaryotic genome annotation. Bioinformatics 30, (2014).

41. Ghaly, T. M., Rajabal, V., Russel, D., Colombi, E. & Tetu, S. G. EcoFoldDB: Protein structure-guided functional profiling of ecologically relevant microbial traits at the metagenome scale. 2025.04.02.646905 Preprint at 10.1101/2025.04.02.646905 (2025).

42. van Kempen, M. et al. Fast and accurate protein structure search with Foldseek. Nat Biotechnol 42, 243–246 (2024).

43. Lombard, V., Golaconda Ramulu, H., Drula, E., Coutinho, P. M. & Henrissat, B. The carbohydrate-active enzymes database (CAZy) in 2013. Nucleic Acids Research 42, (2014).

44. Rice, P., Longden, I. & Bleasby, A. EMBOSS: the European Molecular Biology Open Software Suite. Trends Genet 16, 276–277 (2000).

45. Asnicar, F. et al. Precise phylogenetic analysis of microbial isolates and genomes from metagenomes using PhyloPhlAn 3.0. Nature Communications 11, (2020).

46. R Core Team. R: A Language and Environment for Statistical Computing_. R Foundation for Statistical Computing, Vienna, Austria. (2023).

47. Martínez-Espinosa, R. M. Microorganisms and Their Metabolic Capabilities in the Context of the Biogeochemical Nitrogen Cycle at Extreme Environments. International Journal of Molecular Sciences 21, 4228 (2020).

48. Perez, H. G., Stevenson, C. K., Lourenco, J. M. & Callaway, T. R. Understanding Rumen Microbiology: An Overview. Encyclopedia 4, 148–157 (2024).

49. Jeilu, O., Gessesse, A., Simachew, A., Johansson, E. & Alexandersson, E. Prokaryotic and eukaryotic microbial diversity from three soda lakes in the East African Rift Valley determined by amplicon sequencing. Front Microbiol 13, 999876 (2022).

50. Wu, Y., Jiao, C., Diao, Q. & Tu, Y. Effect of Dietary and Age Changes on Ruminal Microbial Diversity in Holstein Calves. Microorganisms 12, 12 (2023).

51. Huws, S. A. et al. Addressing Global Ruminant Agricultural Challenges Through Understanding the Rumen Microbiome: Past, Present, and Future. Front Microbiol 9, 2161 (2018).

52. Yan, L. et al. Environmental selection shapes the formation of near-surface groundwater microbiomes. Water Research 170, 115341 (2020).

53. Junkins, E. N., McWhirter, J. B., McCall, L.-I. & Stevenson, B. S. Environmental structure impacts microbial composition and secondary metabolism. ISME COMMUN. 2, 1–10 (2022).

54. Gharechahi, J., Sarikhan, S., Han, J.-L., Ding, X.-Z. & Salekdeh, G. H. Functional and phylogenetic analyses of camel rumen microbiota associated with different lignocellulosic substrates. NPJ Biofilms Microbiomes 8, 46 (2022).

55. Stewart, R. D. et al. Compendium of 4,941 rumen metagenome-assembled genomes for rumen microbiome biology and enzyme discovery. Nat Biotechnol 37, 953–961 (2019).

56. Conteville, L. C. et al. Recovery of metagenome-assembled genomes from the rumen and fecal microbiomes of Bos indicus beef cattle. Sci Data 11, 1385 (2024).

57. Arias, P. M. et al. Environment and taxonomy shape the genomic signature of prokaryotic extremophiles. Sci Rep 13, 16105 (2023).

58. Luiselli, J., Rouzaud-Cornabas, J., Lartillot, N. & Beslon, G. Genome Streamlining: Effect of Mutation Rate and Population Size on Genome Size Reduction. Genome Biology and Evolution 16, evae250 (2024).

59. Arella, D., Dilucca, M. & Giansanti, A. Codon usage bias and environmental adaptation in microbial organisms. Mol Genet Genomics 296, 751–762 (2021).

60. Khan, M. F. & Patra, S. Deciphering the rationale behind specific codon usage pattern in extremophiles. Sci Rep 8, 15548 (2018).

61. Chai, J., Zhuang, Y., Cui, K., Bi, Y. & Zhang, N. Metagenomics reveals the temporal dynamics of the rumen resistome and microbiome in goat kids. Microbiome 12, 14 (2024).

62. Pereira, A. M., de Lurdes Nunes Enes Dapkevicius, M. & Borba, A. E. S. Alternative pathways for hydrogen sink originated from the ruminal fermentation of carbohydrates: Which microorganisms are involved in lowering methane emission? Animal Microbiome 4, 5 (2022).

63. Comtet-Marre, S. et al. Metatranscriptomics Reveals the Active Bacterial and Eukaryotic Fibrolytic Communities in the Rumen of Dairy Cow Fed a Mixed Diet. Front. Microbiol. 8, (2017).

64. Takizawa, S. et al. Relationship Between Rumen Microbial Composition and Fibrolytic Isozyme Activity During the Biodegradation of Rice Straw Powder Using Rumen Fluid. Microbes Environ 38, ME23041 (2023).

65. Dao, T.-K. et al. Understanding the Role of Prevotella Genus in the Digestion of Lignocellulose and Other Substrates in Vietnamese Native Goats’ Rumen by Metagenomic Deep Sequencing. Animals (Basel) 11, 3257 (2021).

66. Betancur-Murillo, C. L., Aguilar-Marín, S. B. & Jovel, J. Prevotella: A Key Player in Ruminal Metabolism. Microorganisms 11, 1 (2022).

67. Shinkai, T. et al. Characteristics of rumen microbiota and Prevotella isolates found in high propionate and low methane-producing dairy cows. Front. Microbiol. 15, (2024).

68. Sato, Y. et al. Identification of 146 Metagenome-assembled Genomes from the Rumen Microbiome of Cattle in Japan. Microbes Environ 37, ME22039 (2022).

69. Sato, Y., Sato, R., Fukui, E. & Yoshizawa, F. Impact of rumen microbiome on cattle carcass traits. Sci Rep 14, 6064 (2024).

70. Hernández-Soto, L. M., Martínez-Abarca, F., Ramírez-Saad, H., López-Pérez, M. & Aguirre-Garrido, J. F. Genome analysis of haloalkaline isolates from the soda saline crater lake of Isabel Island; comparative genomics and potential metabolic analysis within the genus Halomonas. BMC Genomics 24, 696 (2023).

71. Pujalte, M. J., Lucena, T., Ruvira, M. A., Arahal, D. R. & Macián, M. C. The Family Rhodobacteraceae. in The Prokaryotes: Alphaproteobacteria and Betaproteobacteria (eds. Rosenberg, E., DeLong, E. F., Lory, S., Stackebrandt, E. & Thompson, F.) 439– 512 (Springer, Berlin, Heidelberg, 2014). doi:10.1007/978-3-642-30197-1_377.

72. Ersoy Omeroglu, E., et al. Microbial community of soda Lake Van as obtained from direct and enriched water, sediment and fish samples. Sci Rep 11, 18364 (2021).

73. Ahn, A.-C. et al. Genomic diversity within the haloalkaliphilic genus Thioalkalivibrio. PLOS ONE 12, e0173517 (2017).

74. Berben, T., Overmars, L., Sorokin, D. Y. & Muyzer, G. Diversity and Distribution of Sulfur Oxidation-Related Genes in Thioalkalivibrio, a Genus of Chemolithoautotrophic and Haloalkaliphilic Sulfur-Oxidizing Bacteria. Front Microbiol 10, 160 (2019).

75. Vavourakis, C. D. et al. Metagenomic Insights into the Uncultured Diversity and Physiology of Microbes in Four Hypersaline Soda Lake Brines. Front Microbiol 7, 211 (2016).

76. Andreote, A. P. D. et al. Contrasting the Genetic Patterns of Microbial Communities in Soda Lakes with and without Cyanobacterial Bloom. Front Microbiol 9, 244 (2018).

77. Stergiadis, S. et al. Unravelling the Role of Rumen Microbial Communities, Genes, and Activities on Milk Fatty Acid Profile Using a Combination of Omics Approaches. Front. Microbiol. 11, (2021).

78. Andersen, T. O. et al. Metabolic influence of core ciliates within the rumen microbiome. ISME J 17, 1128–1140 (2023).

79. Yang, J., Kalhan, S. C. & Hanson, R. W. What Is the Metabolic Role of Phosphoenolpyruvate Carboxykinase? J Biol Chem 284, 27025–27029 (2009).

80. Yu, S., Meng, S., Xiang, M. & Ma, H. Phosphoenolpyruvate carboxykinase in cell metabolism: Roles and mechanisms beyond gluconeogenesis. Mol Metab 53, 101257 (2021).

81. Gharechahi, J. et al. Lignocellulose degradation by rumen bacterial communities: New insights from metagenome analyses. Environmental Research 229, 115925 (2023).

82. Gunde-Cimerman, N., Plemenitaš, A. & Oren, A. Strategies of adaptation of microorganisms of the three domains of life to high salt concentrations. FEMS Microbiology Reviews 42, 353–375 (2018).

83. Jones, D. L. & Baxter, B. K. DNA Repair and Photoprotection: Mechanisms of Overcoming Environmental Ultraviolet Radiation Exposure in Halophilic Archaea. Front. Microbiol. 8, (2017).

84. Sorokin, D. Y. The Microbial Sulfur Cycle at Extremely Haloalkaline Conditions of Soda Lakes. Front. Microbiol. 2, (2011).

85. Li, B. et al. Rumen microbiota of indigenous and introduced ruminants and their adaptation to the Qinghai–Tibetan plateau. Front. Microbiol. 13, (2022).

86. He, B., Jin, S., Cao, J., Mi, L. & Wang, J. Metatranscriptomics of the Hu sheep rumen microbiome reveals novel cellulases. Biotechnology for Biofuels 12, 153 (2019).

87. Neves, A. L. A. et al. Accelerated discovery of novel glycoside hydrolases using targeted functional profiling and selective pressure on the rumen microbiome. Microbiome 9, 229 (2021).

88. Li, J. et al. A catalog of microbial genes from the bovine rumen unveils a specialized and diverse biomass-degrading environment. GigaScience 9, giaa057 (2020).

89. Gao, H. et al. Metagenomic Sequencing Reveals the Taxonomic and Functional Characteristics of Rumen Micro-organisms in Gayals. Microorganisms 11, 1098 (2023).

90. Wang, L., Zhang, G., Xu, H., Xin, H. & Zhang, Y. Metagenomic Analyses of Microbial and Carbohydrate-Active Enzymes in the Rumen of Holstein Cows Fed Different Forage-to-Concentrate Ratios. Front Microbiol 10, 649 (2019).

91. Krause, D. O. et al. Opportunities to improve fiber degradation in the rumen: microbiology, ecology, and genomics. FEMS Microbiology Reviews 27, 663–693 (2003).

92. Sorokin, D. Yu. & Kuenen, J. G. Haloalkaliphilic sulfur-oxidizing bacteria in soda lakes. FEMS Microbiology Reviews 29, 685–702 (2005).

93. Zorz, J. K. et al. A shared core microbiome in soda lakes separated by large distances. Nat Commun 10, 4230 (2019).

94. Parks, D. H. et al. Recovery of nearly 8,000 metagenome-assembled genomes substantially expands the tree of life. Nat Microbiol 2, 1533–1542 (2017).

